# Weakly migratory metastatic breast cancer cells activate fibroblasts via microvesicle-Tg2 to facilitate dissemination and metastasis

**DOI:** 10.1101/2021.10.27.466095

**Authors:** Samantha C. Schwager, Lauren A. Hapach, Caroline M. Carlson, Jenna A. Mosier, Tanner J. McArdle, Wenjun Wang, Anissa L. Jayathilake, Francois Bordeleau, Marc A. Antonyak, Richard A. Cerione, Cynthia A. Reinhart-King

## Abstract

Cancer cell migration is highly heterogeneous, and the migratory capability of cancer cells is thought to be an indicator of metastatic potential. It is becoming clear that a cancer cell does not have to be inherently migratory to metastasize, with weakly migratory cancer cells often found to be highly metastatic. However, the mechanism through which weakly migratory cells escape from the primary tumor remains unclear. Here, utilizing phenotypically sorted highly and weakly migratory breast cancer cells, we demonstrate that weakly migratory metastatic cells disseminate from the primary tumor via communication with stromal cells. While highly migratory cells are capable of single cell migration, weakly migratory cells rely on cell-cell signaling with fibroblasts to escape the primary tumor. Weakly migratory cells release microvesicles rich in tissue transglutaminase 2 (Tg2) which activate fibroblasts and lead weakly migratory cancer cell migration *in vitro*. These microvesicles also induce tumor stiffening and fibroblast activation *in vivo* and enhance the metastasis of weakly migratory cells. Our results identify microvesicles and Tg2 as potential therapeutic targets for metastasis and reveal a novel aspect of the metastatic cascade in which weakly migratory cells release microvesicles which activate fibroblasts to enhance cancer cell dissemination.

## Introduction

It is known that cancer cell migration *in vivo* can be highly heterogeneous with cells exhibiting a wide range of migratory phenotypes, including amoeboid, mesenchymal, single cell, and collective migration(1–3). Even within a single population, cells can exhibit different migration phenotypes(4–7) resulting from intrinsic cancer cell genetic differences(8–11) and extrinsic factors, such as interactions with the extracellular matrix or stromal cells(1,12–17). As the first step in the metastatic cascade involves the migration and invasion of cancer cells away from the primary tumor, migratory capability is largely believed to be an indicator of cancer progression(18–21). However, it is now becoming clear that a cancer cell does not have to be inherently migratory to metastasize(22–27). In fact, some of the highly metastatic phenotypes in breast and colorectal cancers are often less migratory than the weakly metastatic phenotypes, yet they are still able to enter the circulation as efficiently(23,24,26–28). While enhanced clustering, survival, and proliferation have been suggested as potential mechanisms for why weakly migratory cells can outperform their highly migratory counterparts in the late stages of the metastatic cascade thus contributing to metastasis(24,26), it is not clear how these weakly migratory cells can efficiently escape from the primary site.

To escape the primary site during local invasion, cancer cells navigate through a heterogeneous tumor microenvironment, where they interact with the extracellular matrix and a diverse collection of stromal cells. Chemical, physical, and metabolic interactions with the tumor microenvironment are known to alter the invasion capacity of cancer cells(14,29,30). Cancer associated fibroblasts (CAFs) are a stromal cell in the tumor microenvironment that can lead and promote tumor cell invasion and metastasis through extracellular matrix (ECM) remodeling(14,15,31–35). CAFs at the primary tumor are largely derived from fibroblasts that have been transformed to a more contractile, activated state(36), and cancer-derived microvesicles (MVs) have recently been implicated in fibroblast activation(37–39). Thus, we hypothesized that while highly migratory cells can escape the primary tumor independently, CAFs facilitate weakly migratory cancer cell escape from the primary tumor.

Here, we identify a novel aspect of the metastatic cascade by which weakly migratory cancer cells release tissue transglutaminase 2 (Tg2)-rich MVs to activate fibroblasts and enhance cancer cell dissemination. Moreover, highly and weakly migratory cells release MVs which differentially signal to the tumor microenvironment. MVs from highly migratory cells have little effect on fibroblast activation, and highly migratory cells do not require activated fibroblasts to migrate. In contrast, MVs from weakly migratory cells are rich in tissue transglutaminase 2 (Tg2) and activate fibroblasts which enhance fibroblast-led weakly cancer cell migration *in vitro*. These MVs also induce tumor stiffening and fibroblast activation *in vivo* and enhance the metastasis of weakly migratory cells. Our findings highlight MVs and Tg2 as potential targets for developing therapeutics to prevent metastasis.

## Results

### Highly and weakly migratory breast cancer subpopulations form tumors with distinct matrix and fibroblast populations

To investigate the mechanism through which weakly migratory cells locally invade and disseminate, MDA-MB-231 breast cancer cells were phenotypically sorted based on migration through a collagen coated transwell(4). Twenty rounds of sorting resulted in the isolation of two stable subpopulations of breast cancer cells: highly migratory MDA^+^ and weakly migratory MDA^-^ (Fig. 1a). Consistent with our previous findings(4), the highly migratory MDA^+^ subpopulation was weakly metastatic in an orthotopic mouse model while the weakly migratory MDA^-^ subpopulation was highly metastatic (Fig. 1b-c). Given the differing migratory and metastatic capabilities of MDA^+^ and MDA^-^, we sought to characterize the tumor microenvironment formed by highly and weakly migratory cell *in vivo* to determine the impact of the tumor microenvironment on cancer cell dissemination and subsequent metastasis.

**Figure 1.**
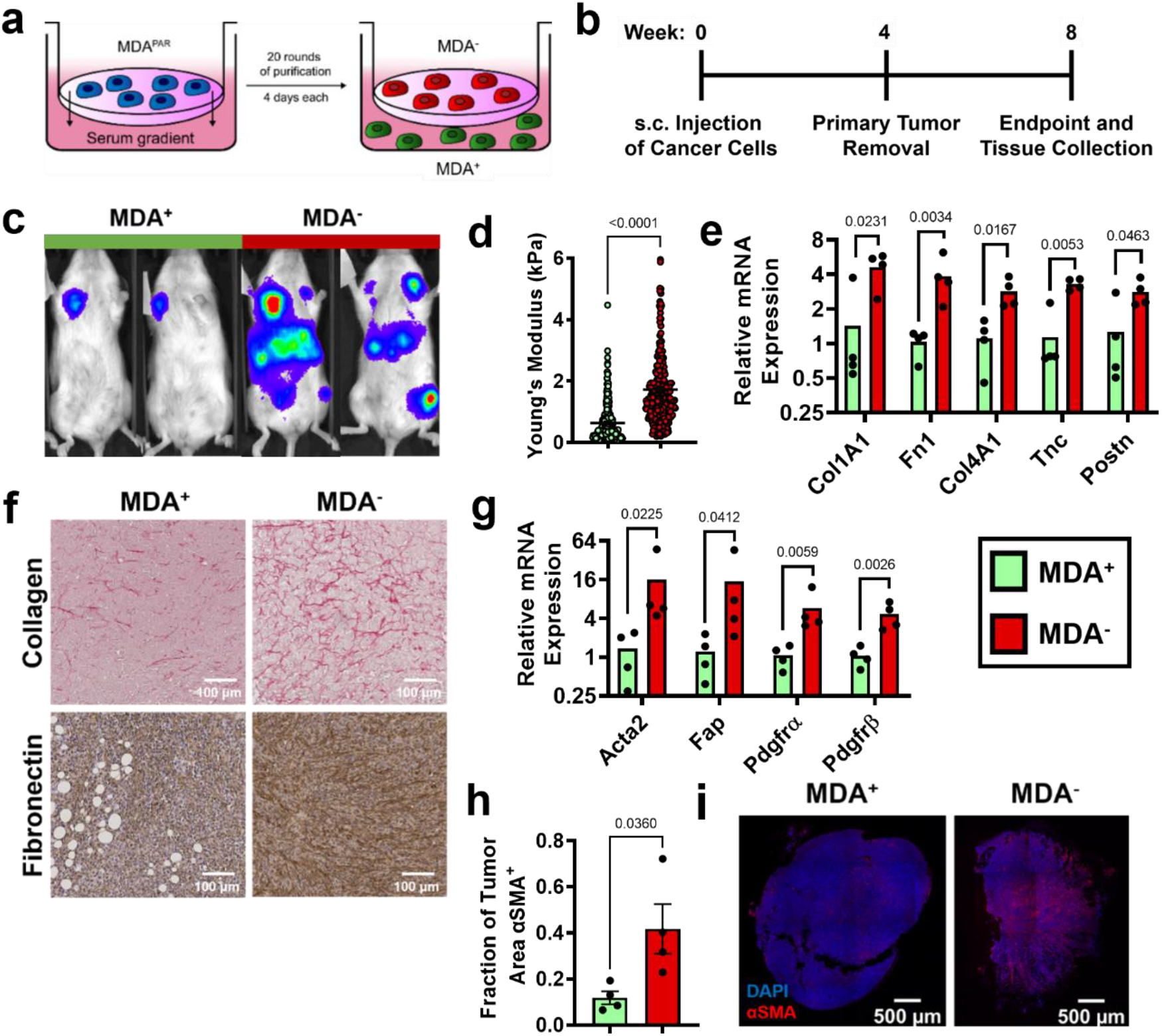
Highly and weakly migratory breast cancer subpopulations form tumors with distinct matrix and fibroblast populations. (a) Schematic of phenotypic sorting. (b) Timeline of orthotopic mammary metastasis model. (c) Representative BLI of MDA^+^ and MDA^-^ metastasis. (d) AFM stiffness measurements of MDA^+^ and MDA^-^ primary tumors. (N=3+ tumors per condition; n = 254, 280). (e) Relative mRNA expression of mouse-derived matrix components in MDA^+^ and MDA^-^ primary tumors. (N=4 tumors per condition). (f) Immunohistochemical staining of collagen and fibronectin in MDA^+^ and MDA^-^ primary tumors. (g) Relative mRNA expression of stromal CAF markers in MDA^+^ and MDA^-^. (N=4 tumors per condition). (h) Fraction of tumor area positive for αSMA (N=4). (e) Representative images of αSMA (red) and DAPI (blue) in MDA^+^ and MDA^-^ tumors. mRNA graphs show mean + individual data points. Bar graphs shown mean +/− SEM. P-values determined using an unpaired Student’s t-test.

Since increased collagen and fibronectin deposition and increased tumor stiffness have been linked to tumor progression(40–42), we investigated the mechanical properties of the MDA^+^ and MDA^-^ tumors to assess whether altered tumor matrix mechanics may contribute to metastatic potential. Based on Atomic Force Microscopy measurements, MDA^-^ primary tumors exhibited significantly higher tumor stiffness compared to MDA^+^ tumors (Fig. 1d). qPCR of tumors revealed significantly higher amounts of mouse-derived matrix, including collagen I (Col1A1), fibronectin 1 (Fn1), collagen IV (Col4A1), tenascin-C (Tnc), and periostin (Postn), in MDA^-^ tumors compared to MDA^+^ tumors (Fig. 1e). Similarly, increased collagen and fibronectin was evident via immunohistochemical staining in MDA^-^ tumors compared to MDA^+^ tumors (Fig. 1f). These results indicate that the weakly migratory, highly metastatic MDA^-^ form stiffer tumors with increased ECM.

Given that our data indicate MDA^-^ produce stiffer tumors than MDA^+^, and CAFs are a major stromal component of breast tumors that mediate matrix deposition(43), we investigated the CAF component of tumors from each subpopulation. MDA^-^ tumors exhibited increased levels of several CAF markers, including α-smooth muscle actin (Acta2), fibroblast activation protein (Fap), and platelet-derived growth factors α and β (Pdgfrα, Pdgfrβ) (Fig. 1g). MDA^-^ tumors also exhibited an increased fraction of αSMA positive tissue area compared to MDA^+^ tumors (Fig. 6h-i). These findings reveal that the weakly migratory, highly metastatic MDA^-^ primary tumors have increased fibroblast activation.

### MVs released from weakly migratory cancer cells are potent activators of fibroblasts in vitro

Since our data indicate that MDA^-^ tumors are stiffer and have a larger population of CAFs, and it is known that MVs can induce fibroblast activation(37,38), we compared the MVs released from MDA^+^ and MDA^-^. MVs were collected from the MDA^+^ and MDA^-^ subpopulations using size-based filtration, termed MV^+^ and MV^-^, respectively. MDA^-^ released significantly more MVs than MDA^+^ (Fig. 2a) but both MV^+^ and MV^-^ had similar size distributions (Fig. 2b). To probe MV signaling to fibroblasts, fibroblast phenotypes associated with fibroblast activation and cancer progression were examined after culture with MV^+^ and MV^-^ on 20 kPa polyacrylamide gels, representing the stiffness of breast ECM at the tumor periphery(44) (Fig. 2c). MV^-^ caused focal adhesion kinase (FAK) Tyr397 phosphorylation in fibroblasts, as increased FAK phosphorylation (pFAK) was evident in fibroblasts cultured with MV^-^ compared to control and MV^+^ conditions (Fig. 2d), consistent with previous findings that MDA-MB-231 MVs regulate fibroblast FAK activation(37). Additionally, fibroblasts cultured with MV^-^ exhibited increased α-smooth muscle actin (αSMA) expression, a marker of fibroblast activation, and cell area compared to both control and MV^+^ conditions (Fig. 2e-f). Fibroblasts cultured with MV^+^ displayed no change in αSMA expression or cell area compared to control conditions (Fig. 2e-f). Fibroblasts cultured with MV^-^ also displayed increased positive fibronectin staining (Fig. 2g, Supp. Fig. 1b), suggesting the MV^-^ increased fibronectin deposition by fibroblasts. Fibroblasts cultured with MV^-^ exhibited increased EdU incorporation compared to MV^+^ and control conditions, indicative of increased proliferation (Fig. 2h, Supp. Fig. 1a). Increased traction force and traction stress was evident in fibroblasts cultured with MV^-^, compared to control conditions, but not in fibroblasts cultured with MV^+^ (Fig. 2i, Supp. Fig. 1d-e). Overall, this data indicates that MV^-^ activate fibroblasts to a more proliferative, contractile state compared to MV^+^ and control conditions.

**Figure 2.**
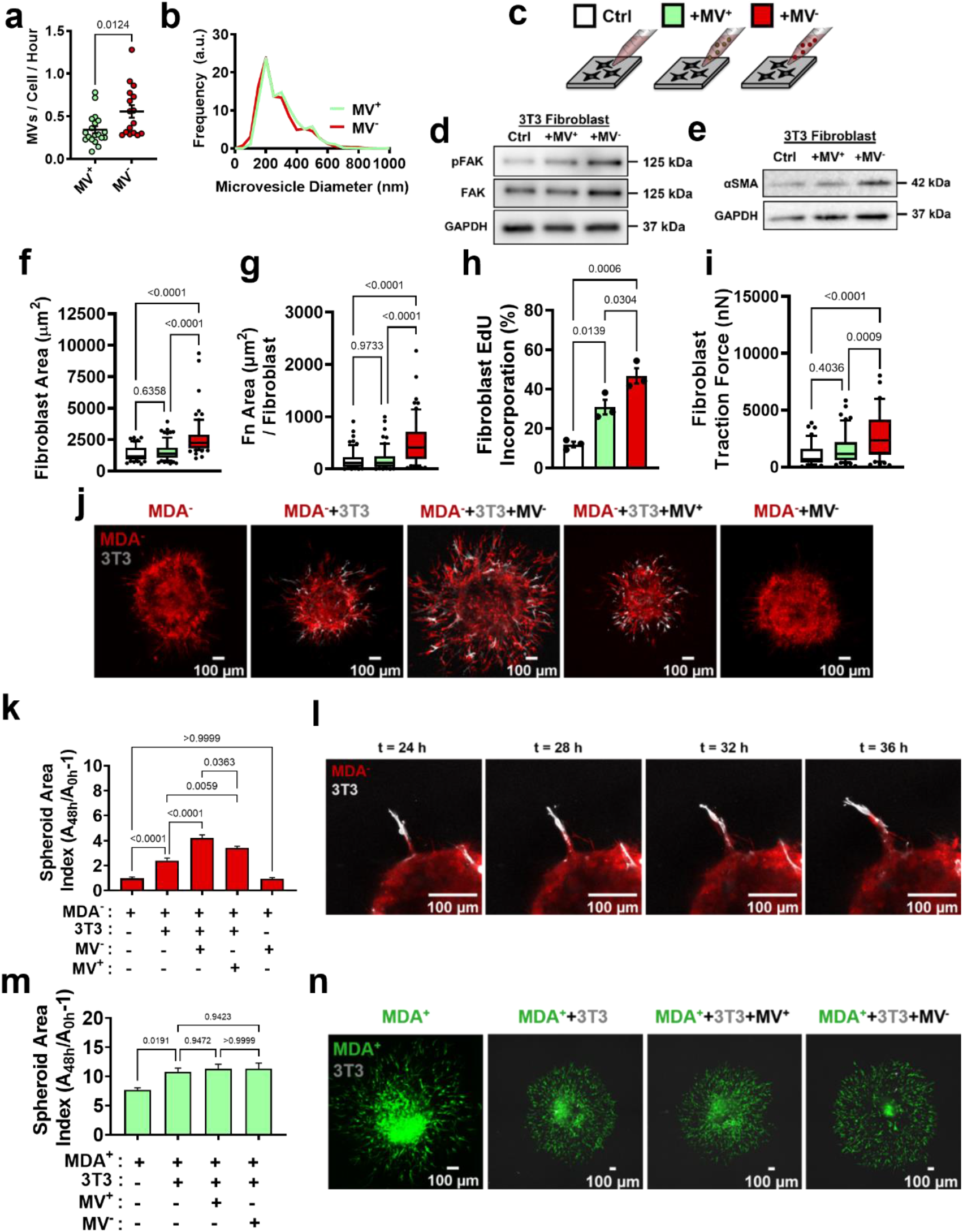
MV^-^ are potent activators of fibroblasts *in vitro*. (a) Number of MVs released from MDA^+^ and MDA^-^ per hour (N=19, 16). (b) Size distribution of MVs isolated from MDA^+^ and MDA^-^. (c) Schematic overviewing fibroblast culture with MVs. (d) Western blot of pFAK, FAK, and GAPDH in fibroblasts cultured in cultured in control conditions (Ctrl), with MV^+^ (+MV^+^), or with MV^-^ (+MV^-^). (e) Western blot of αSMA and GAPDH in fibroblasts cultured in in Ctrl, +MV^+^, or +MV^-^ conditions (f) Fibroblast cell area after culture in Ctrl, +MV^+^, or +MV^-^ conditions. (N=3; n=69, 75, 77). (g) Area of fibronectin deposited per fibroblast cultured in Ctrl, +MV^+^, or +MV^-^ conditions. (N=3; n=66, 66, 61). (h) Percentage of fibroblasts EdU positive after culture in Ctrl, +MV^+^, or +MV^-^ conditions. (N=3). (i) Fibroblast traction force after culture in Ctrl, +MV^+^, or +MV^-^ conditions. (N=3; n=41, 57, 58). (j) Representative images of spheroid outgrowth 48 hours post embedding. MDA^-^ (red); 3T3 fibroblast (gray). (k) Spheroid area index 48 hours post embedding. (N=3+; n=22, 31, 32, 22, 19). (l) Time series images of fibroblast leading MDA^-^ escape from spheroid. (m) Spheroid area index 48 hours post embedding. (N=3+; n=28, 62, 34, 28). (n) Representative images of spheroid outgrowth 48 hours post embedding. MDA^+^ (green); 3T3 fibroblast (gray). Bar graphs show mean +/− SEM. Box and whisker plots show median and 25^th^-75^th^ (box) and 10^th^-90^th^ (whiskers) percentiles. P-values determined using an unpaired Student’s t-test or a one-way ANOVA with Tukey’s test for multiple comparisons.

### MV-mediated fibroblast activation increases weakly migratory cancer cell migration

CAFs enable cancer cell migration and metastasis, in part, through matrix remodeling at the primary tumor(14,15,31–35). Given that our data indicate that MDA^-^ cells induce fibroblast activation through the release of MV, we investigated whether MV-activated fibroblasts could promote the migration of MDA^-^ in a tumor spheroid model. After 48 hours of culture, MDA^-^ spheroids exhibited minimal spheroid outgrowth, measured using a spheroid area index (A_48h_/A_0h-1_) (Fig. 2i-j, Supp. Fig. 1f-g), while MDA^+^ spheroids exhibited high levels of spheroid outgrowth (Fig. 2m-n, Supp. Fig. 1f,i), consistent with our previous findings(4). Coculture of MDA^-^ or MDA^+^ with 3T3 fibroblasts (MDA^-^+3T3, MDA^+^+3T3) resulted in significantly enhanced spheroid outgrowth compared to cancer cells alone (Fig. 2j-k, m-n, Supp. Fig. 1f-g, i). MDA^-^+3T3 spheroids displayed fibroblast-led strands of MDA^-^ migration away from the spheroid, suggesting that the fibroblasts physically lead the migration of MDA^-^ through the matrix (Fig. 2l). Importantly, MDA^-^+3T3 spheroids cultured with MV^-^ exhibited significantly enhanced spheroid outgrowth and MDA^-^ migration distance, compared to MDA^-^+3T3 control spheroids (Fig. 2j-k, Supp. Fig. 1f-g). MDA^-^+3T3 spheroids cultured with MV^+^ also showed increased outgrowth compared to MDA^-^+3T3 spheroids, but this outgrowth was significantly less than that induced by MV^-^ (Fig. 2j-k, Supp. Fig. 1f-g). Interestingly, MDA^+^+3T3 spheroids cultured with MV^+^ or MV^-^ exhibited no change in spheroid outgrowth and MDA^+^ migration distance compared to control conditions (Fig. 2m-n, Supp. Fig. 1f,i). This result suggests that MDA^+^ are capable of robust migration independently of MV^-^mediated fibroblast signaling. When MV^-^ were applied to MDA^-^ only spheroids, no increase in spheroid outgrowth was observed, highlighting the necessity of fibroblasts for this MV^-^induced cancer cell migration (Fig. 2j-k, Supp. Fig. 1f-g). Additionally, MDA^-^ spheroids cultured with fibroblast conditioned media or MV^-^-treated fibroblast conditioned media displayed no change in spheroid outgrowth (Supp. Fig. 1h), indicating that fibroblast-enhanced cancer cell migration is not a result of secreted factors but rather due to physical interactions. Altogether, these results indicate that MV^-^ are potent activators of fibroblasts and that MV^-^-induced fibroblast activation mechanically feeds back to induce the migration of MDA^-^.

### Highly and weakly migratory breast cancer cells release MVs with distinctly different contents

To determine the MV cargo responsible for the differential activation of fibroblasts by the MDA^-^ cells, iTRAQ proteomics was completed on MV^+^ and MV^-^. Identified proteins were compared to Exocarta and Vesiclepedia EV databases and exhibited significant overlap with both EV databases (Fig. 3a). Proteomics data suggests that MV cargo was different between highly migratory and weakly migratory subpopulations of breast cancer cells, with 26.8% of the identified proteins being more highly expressed by MV^+^ (FC≥1.1) and 29.8% being more highly expressed by MV^-^ (FC≤0.9) (Fig. 3b-c). 43.4% of identified proteins were similarly expressed between both MV populations (0.9<FC<1.1) (Fig. 3b-c). Protein set enrichment analysis (PSEA) of MV^+^ and MV^-^ cargo was performed using the PSEA-Quant algorithm(45). PSEA revealed different GO annotations for MV^+^ and MV^-^ protein expression (Fig. 3d-e). Of note, MV^+^ were enriched for cargo involved in biological adhesion, cell adhesion, location, and cell motility (Fig. 3d). MV^-^ were enriched for cargo involved in various metabolic and catabolic processes and vesicle-mediated transport (Fig. 3e). These findings indicate that cancer cells with varying migratory ability release MVs with distinctly different cargo.

**Figure 3.**
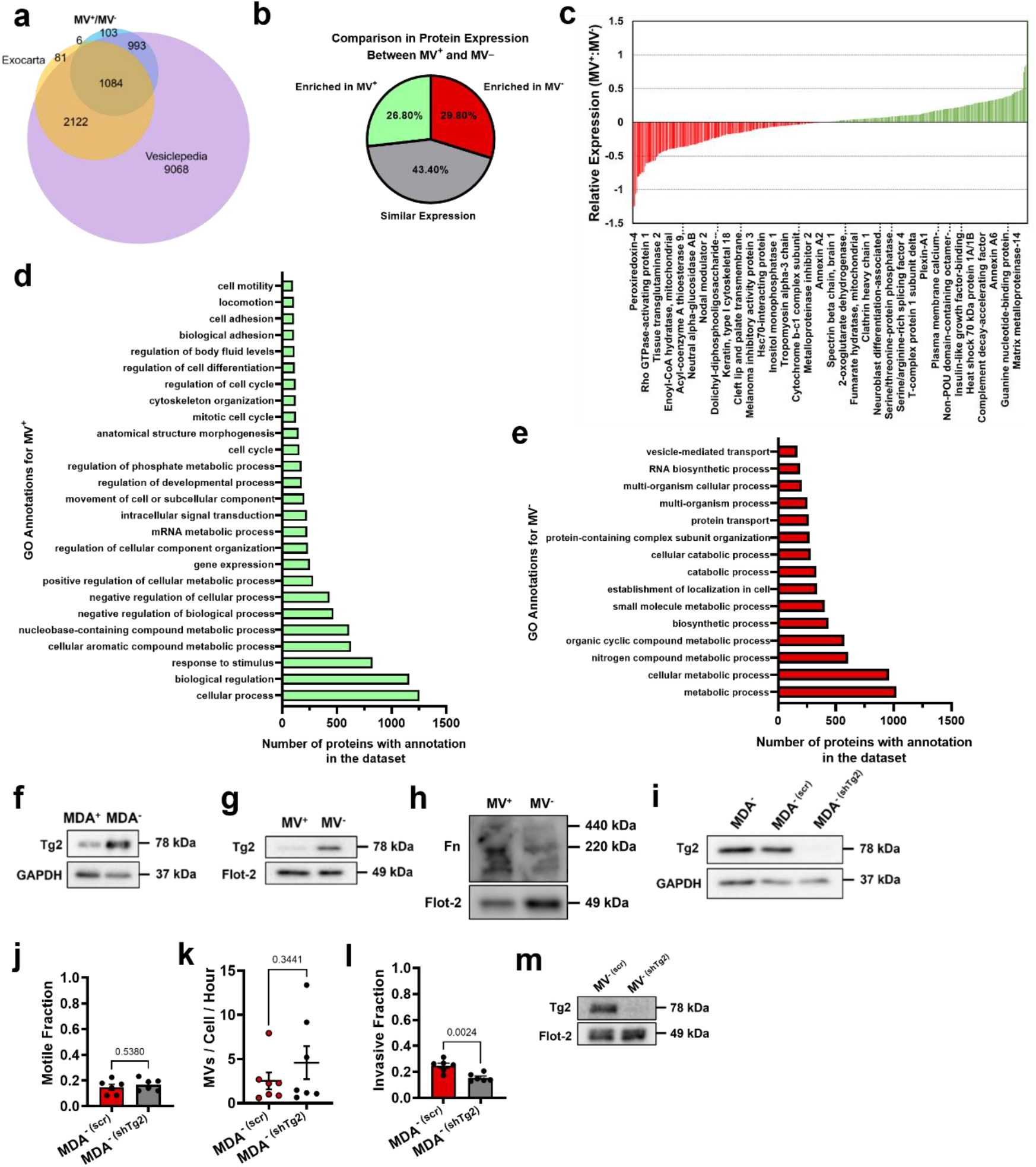
Phenotypically sorted breast cancer subpopulations release MVs with distinctly different contents. (a) Comparison of proteomics of MV^+^ and MV^-^ to Exocarta and Vesiclepedia databases. (b) Comparison of protein expression between MV^+^ and MV^-^. Enriched in MV^+^ = FC≥1.1; similar expression = 0.9<FC<1.1; enriched in MV^-^ = FC≤0.9. (c) Relative expression of proteins identified in proteomics of MV^+^ and MV^-^. Positive values in green are enriched in MV^+^. Negative values in red are enriched in MV^-^. (d) GO annotations for proteins enriched in MV^+^. (e) GO annotations for proteins enriched in MV^-^. (f) Western blot of Tg2 and GAPDH in MDA^+^ and MDA^-^. (g) Western blot of Tg2 and Flot-2 in MV^+^ and MV^-^. (h) Western blot of Fn and Flot-2 in MV^+^ and MV^-^. (i) Western blot of Tg2 and GAPDH in MDA^-^, MDA^- (scr)^, and MDA^- (shTg2)^. (j) Motile fraction of MDA^- (scr)^ and MDA^- (shTg2)^ in 3D collagen. (N=3; n=6). (k) Number of MVs released from MDA^- (scr)^ and MDA^- (shTg2)^ per hour. (N=7). (l) Invasive fraction of MDA^- (scr)^ and MDA^- (shTg2)^ through collagen coated transwells. (N=3; n=6). (m) Western blot of Tg2 and Flot-2 in MV^- (scr)^ and MV^- (shTg2)^. Bar graphs shown mean +/− SEM. P-values determined using an unpaired Student’s t-test.

One of the proteins differentially packaged in MV^-^ compared to MV^+^ was Tg2, a calcium-dependent enzyme that crosslinks collagen and activates fibroblasts (Fig. 3c). EVs have previously been shown to transfer Tg2 to fibroblasts resulting in fibroblast activation(37,46). More specifically, it has been shown that EV-Tg2 crosslinks EV-fibronectin (Fn) into dimers to potentiate fibroblast integrin signaling upon EV-fibroblast interactions(37,46). Increased Tg2 expression in MDA^-^ and MV^-^, compared to MDA^+^ and MV^+^, was verified using western blot (Fig. 3f-g). While the Fn dimer was present in both MV^+^ and MV^-^, it was more highly expressed in MV^+^ (Fig. 3h), suggesting that fibroblast activation by MV^-^ does not solely rely on Fn dimer-mediated integrin signaling.

### Modulation of MV-Tg2 expression regulates MV-mediated fibroblast activation

Given that Tg2 is a known mediator of MV-induced fibroblast transformation(37), we investigated whether knockdown of Tg2 abrogates MV^-^-induced fibroblast activation. MDA^-^ was stably transduced with shRNA targeting Tg2 (MDA^- (shTg2)^) (Fig. 3i). Knockdown of Tg2 resulted in no change in MDA^-^ migration or MV release compared to the scrambled control (MDA^- (scr)^) (Fig. 3j-k). A slight decrease in Transwell invasion was observed in MDA^- (shTg2)^ compared to MDA^- (scr)^ (Fig. 3l). MVs released from MDA^- (shTg2)^ (MV^-shTg2)^) had reduced Tg2 expression compared to the scrambled control MVs (MV^- (scr)^) (Fig. 3m). Fibroblasts cultured with MV^- (shTg2)^ exhibited decreased αSMA expression, cell area, fibronectin area, proliferation, and traction force, compared to culture with MV^- (scr)^ (Fig. 4a-f, Supp. Fig. 2a-e). This suggests that knockdown of Tg2 in MDA-cells reduced the capacity of MV^-^ to activate fibroblasts. Treatment of MV^-^ with the Tg2 inhibitor, T101, decreased fibroblast αSMA expression compared to culture with MV^-^, further supporting role of MV-Tg2 in fibroblast activation (Fig. 4g).

**Figure 4.**
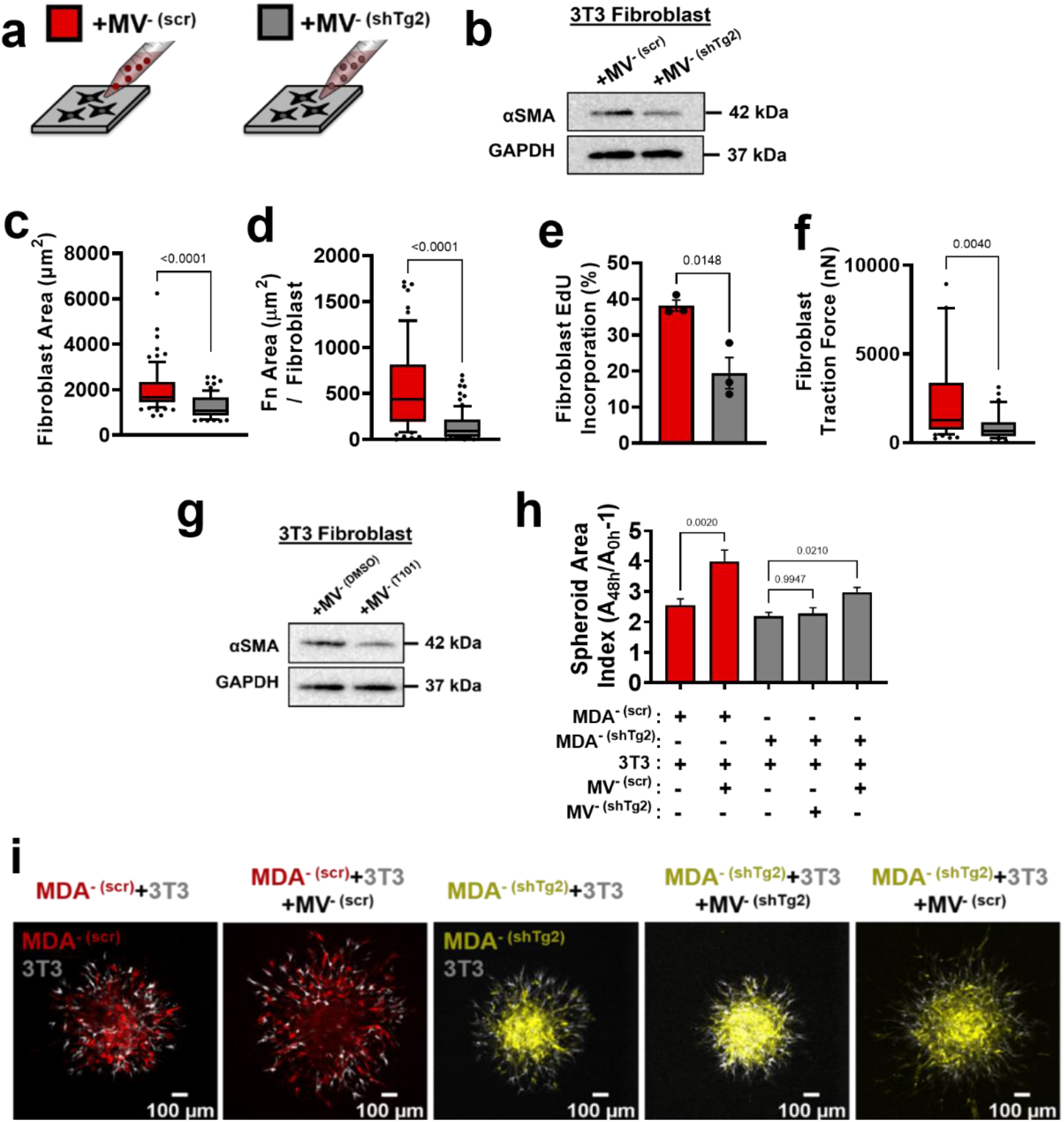
Modulation of Tg2 expression in MVs regulates MV-mediated fibroblast activation. (a) Schematic overviewing fibroblast culture with MVs. (b) Western blot of αSMA and GAPDH in fibroblasts cultured with MV^- (scr)^ (+MV^- (scr)^) or with MV^- (shTg2)^ (+MV^- (shTg2)^). (c) Fibroblast cell area after culture in +MV^- (scr)^ or +MV^- (shTg2)^ conditions. (N=3; n=75,69). (d) Area of fibronectin deposited per fibroblast cultured in +MV^- (scr)^ or +MV^- (shTg2)^ conditions. (N=3; n=62, 82). (e) Percentage of fibroblasts EdU positive after culture in +MV^- (scr)^ or +MV^- (shTg2)^ conditions. (N=3). (f) Fibroblast traction force after culture in +MV^- (scr)^ or +MV^- (shTg2)^ conditions. (N=3; n=56, 33). (g) Western of blot of αSMA and GAPDH in fibroblasts cultured with MV^- (DMSO)^ (+MV^- (DMSO)^) or with MV^- (T101)^ (+MV^- T101^)). (h) Spheroid area index quantification 48 hours post embedding. (N=3+; n=23, 23, 83, 60, 37). (i) Representative images of spheroid outgrowth 48 hours post embedding. MDA^- (scr)^ (red); 3T3 fibroblast (gray); MDA^- (shTg2)^ (yellow). Bar graphs show mean +/− SEM. Box and whisker plots show median and 25^th^-75^th^ (box) and 10^th^-90^th^ (whiskers) percentiles. P-values determined using an unpaired Student’s t-test or a one-way ANOVA with Tukey’s test for multiple comparisons.

Spheroid cocultures of MDA^- (scr)^ and MDA^- (shTg2)^ with 3T3 fibroblasts were utilized to assess the effects of MV^- (shTg2)^ on cell migration. MDA^- (scr)^+3T3 spheroids cultured with MV^- (scr)^ exhibited a significant increase in spheroid outgrowth compared to untreated conditions, consistent with our prior experiments (Fig. 4h-I, Supp. Fig. 2f). MDA^-^ (s^hTg2)^_+_3T3 spheroids cultured with MV^- (shTg2)^ exhibited no change in spheroid outgrowth compared to untreated control conditions (Fig. 4h-I, Supp. Fig. 2f), suggesting that without MV-induced fibroblast activation, fibroblasts could not increase cancer cell migration. MDA^- (shTg2)^+3T3 spheroids cultured with MV^- (scr)^ exhibited a significant increase in spheroid migration compared to untreated conditions, highlighting that MV-Tg2 is required for MV-mediated fibroblast-induced cancer cell migration (Fig. 4h-I, Supp. Fig. 2f).

To further investigate the role of Tg2 in MV-mediated fibroblast activation, Tg2 was overexpressed in MDA^+^ using lentiviral transduction (MDA^+ (FUW-Tg2)^) (Supp. Fig. 2g). MVs released from MDA^+ (FUW-Tg2)^ (MV^+ (FUW-Tg2)^) were confirmed to express increased levels of Tg2 using western blot (Supp. Fig. 2g). Fibroblasts cultured with MV^+ (FUW-Tg2)^ displayed increased cell area, compared to culture with MV^+^ (Supp. Fig. 2i). These results were consistent with earlier findings suggesting that MV-Tg2 regulates MV-mediated fibroblast phenotype (Fig. 4c).

### Tg2 knockdown in MDA^-^ reduces metastasis

Given that MV-Tg2 activated fibroblasts *in vitro* and increased cancer cell migration in spheroid cocultures, we investigated whether knockdown of Tg2 in MDA^-^ is sufficient to reduce MDA^-^ metastasis in an orthotopic mammary metastasis mouse model (Fig. 5a). Six weeks post subcutaneous injection of MDA^- (scr)^ and MDA^- (shTg2)^ into the mammary fat pad, primary tumors were of similar volumes (Fig. 5b). MDA^- (shTg2)^ tumors exhibited decreased stiffness compared to control tumors (Fig. 5c). Immunofluorescent staining of MDA^- (scr)^ and MDA^- (shTg2)^ tumors for the fibroblast activation marker αSMA revealed a decreased fraction of αSMA^+^ tissue in MDA^- (shTg2)^ tumors compared to control tumors (Fig. 5d-e), suggesting that knockdown of Tg2 reduced primary tumor fibroblast activation. Immunohistochemical staining for collagen and fibronectin also revealed decreased matrix in the MDA^- (shTg2)^ tumors compared to control tumors (Fig. 5f). Three weeks post primary tumor removal, BLI imaging and tissue collection was completed. The metastasis of MDA^- (shTg2)^ to the lungs and liver was greatly reduced compared to MDA^- (scr)^ (Fig. 5g-k). Livers from MDA^- (shTg2)^ mice displayed fewer macroscopic liver nodules compared to MDA^- (scr)^ (Fig. 5f), and immunohistochemical staining for GFP^+^ cancer cells in mouse liver and lungs revealed decreased GFP+ tissue area in the lungs and liver of MDA^- (shTg2)^ mice (Fig. 5g-k). These findings reveal that knockdown of Tg2 in MDA^-^ decreases tumor stiffness, fibroblast activation, ECM deposition, and metastasis.

**Figure 5.**
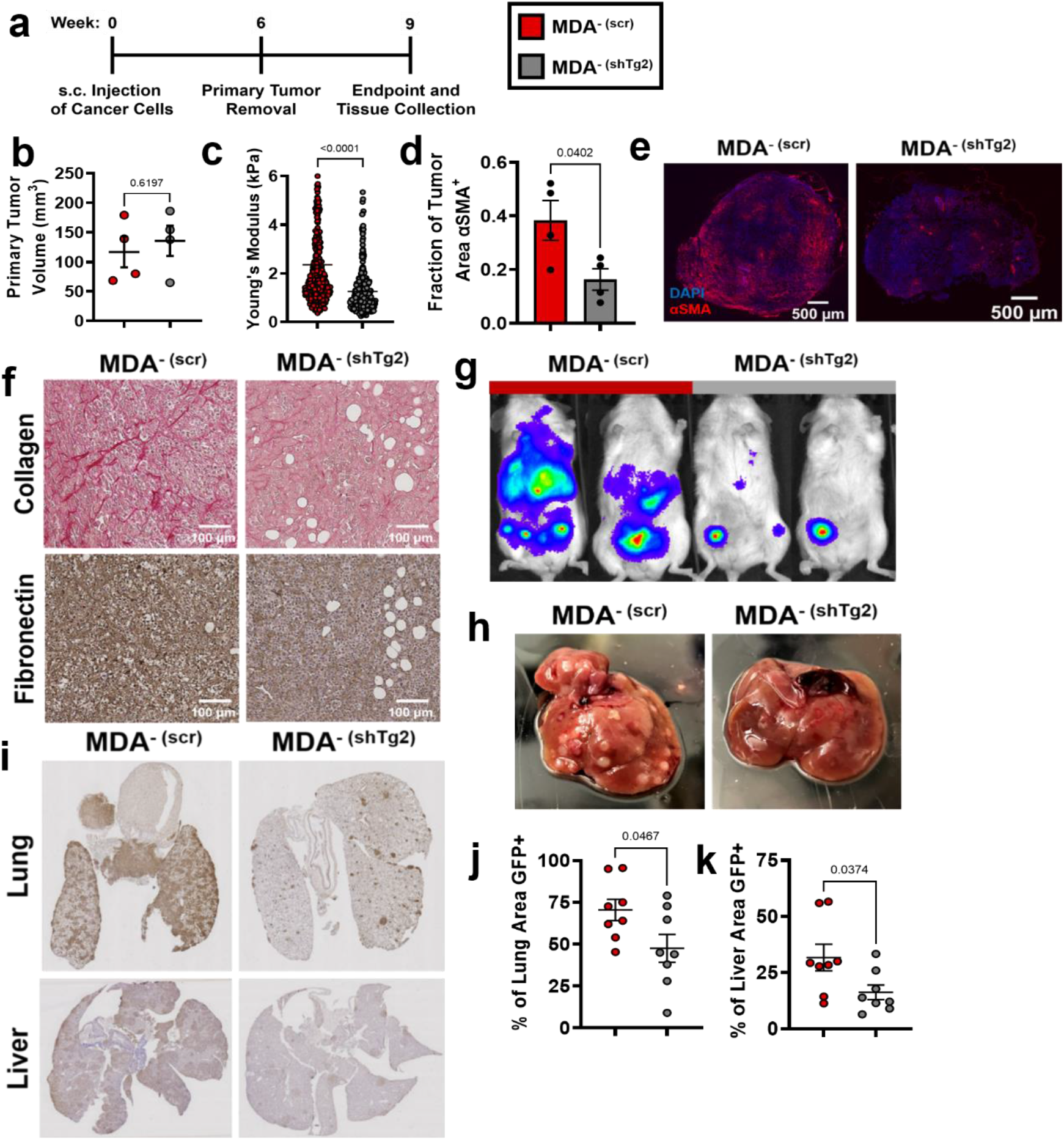
Knockdown of Tg2 in MDA^-^ subpopulation reduces metastasis. (a) Timeline of orthotopic mammary metastasis model. (b) Primary tumor volume after removal (N=4). (c) AFM stiffness measurements of MDA^- (scr)^ and MDA^- (shTg2)^ primary tumors. (N=3+ tumors per condition; n = 420, 379). (d) Fraction of tumor area positive for αSMA (N=4). (e) Representative images of αSMA (red) and DAPI (blue) in MDA^- (scr)^ and MDA^- (shTg2)^ primary tumors. (f) Immunohistochemical staining of collagen and fibronectin in MDA^- (scr)^ and MDA^- (shTg2)^ primary tumors. (g) Representative BLI of MDA^- (scr)^ and MDA^- (shTg2)^ metastasis. (h) Representative image of macroscopic liver nodules. (i) Anti-GFP immunohistochemical staining of lungs and liver from MDA^- (scr)^ and MDA^- (shTg2)^ mice. (j) Quantification of percentage of lung tissue area GFP-positive. (N=4, n=8). (k) Quantification of percentage of liver tissue area GFP-positive. (N=4, n=8). Data shown as mean +/− SEM. P-values determined using an unpaired Student’s t-test.

### Tg2-rich MV^-^ are sufficient to induce MDA^- (shTg2)^ metastasis

Since knockdown of Tg2 in MDA^-^ reduced MDA^-^ metastasis, we investigated whether supplementing primary tumors composed of MDA^- (shTg2)^ cells with Tg2-rich MV^-^ could induce the metastasis of MDA^- (shTg2)^. One week post-subcutaneous injection of MDA^- (shTg2)^ into the mammary fat pad, mice were subcutaneously injected every three days for five weeks with either serum-free (SF) media (MDA^- (shTg2)^ + SF) or MV^-^ (MDA^- (shTg2)^ _+_ MV^-^) (Fig. 6a). After six weeks of primary tumor growth, MDA^- (shTg2)^ primary tumors supplemented with MV^-^ were significantly larger than and exhibited increased primary tumor stiffness compared to control tumors (Fig. 6b-c). MDA^- (shTg2)^ + MV^-^ tumors also exhibited an increased fraction of αSMA^+^ tissue area compared to control tumors (Fig. 6d-e). Immunohistochemical staining of tumors revealed increased collagen and fibronectin deposition in the MDA^- (shTg2)^ + MV^-^ primary tumors compared to control tumors (Fig. 6f). Three weeks post primary tumor removal, BLI imaging and tissue collection was completed. Supplementing MDA^- (shTg2)^ primary tumors with MV^-^ induced increased metastasis to both the lungs and liver of mice, compared to control mice (Fig. 6g-k). Livers from MDA^- (shTg2)^ + MV^-^ mice displayed increased numbers of macroscopic liver nodules compared to control mice (Fig. 6h). Immunohistochemical staining for GFP in mouse lungs revealed increased metastasis to the lungs of MDA^- (shTg2)^ + MV^-^ mice compared to control mice (Fig. 6i-j). Livers of MDA^- (shTg2)^ + MV^-^ mice had slightly increased but not significantly higher levels of metastasis compared to control mice (Fig. 6i,k). These results indicate that repeated injection of Tg2-rich wildtype MV^-^ during the growth of MDA^- (shTg2)^ primary tumors resulted in primary tumor stiffening, increased fibroblast activation, and increased metastasis of the Tg2-knockdown weakly migratory breast cancer subpopulation. Altogether, our results reveal a novel mechanism by which weakly migratory cancer cells release Tg2-rich MVs to activate fibroblasts and remodel the primary tumor to facilitate weakly migratory cancer cell migration and metastasis (Fig. 6l).

**Figure 6.**
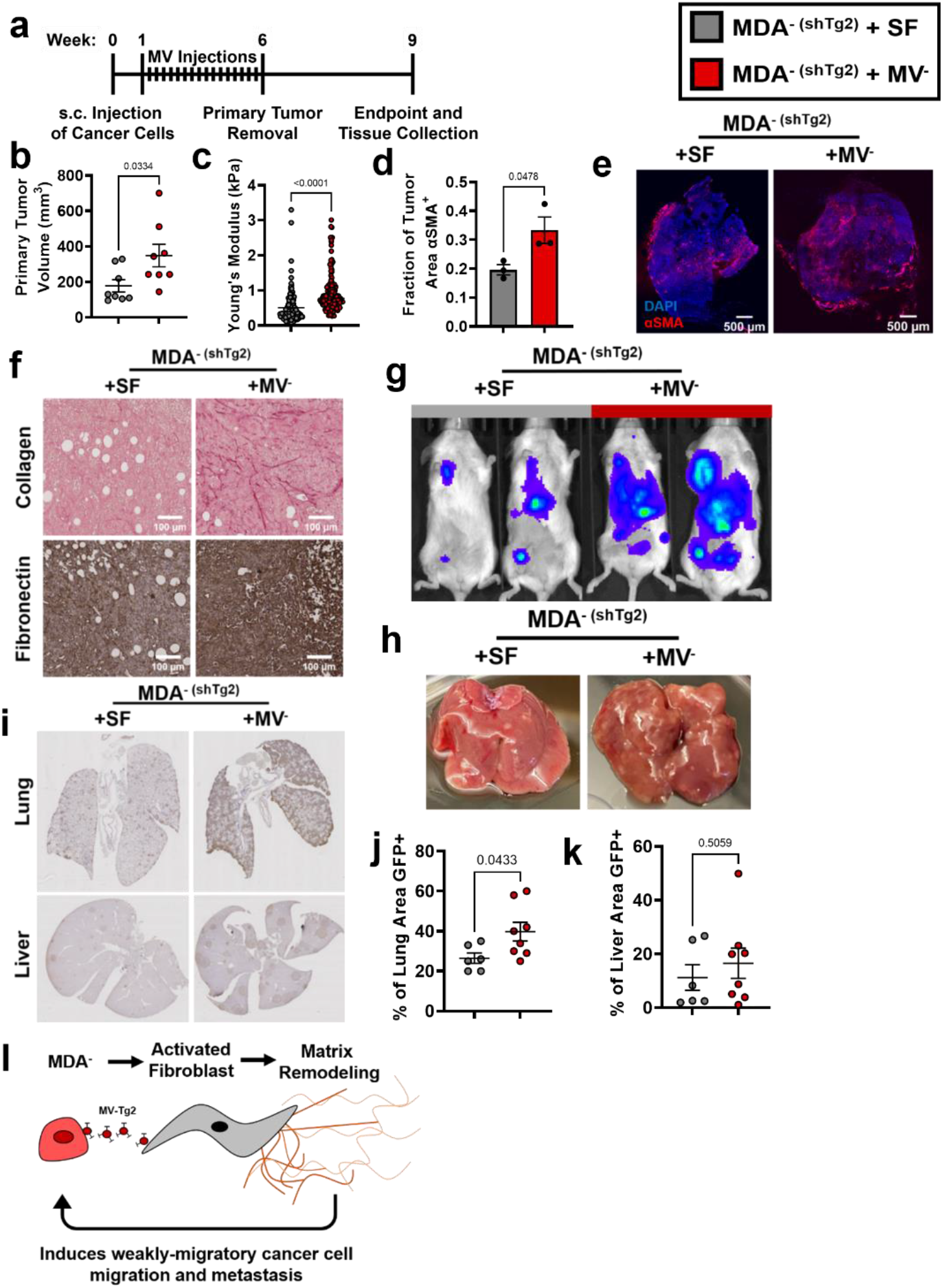
Tg2-rich wildtype MV^-^ are sufficient to induce the metastasis of MDA^- (shTg2)^. (a) Timeline of orthotopic mammary metastasis model with MV injections every 3 days. (b) Primary tumor volume after removal (N=8). (c) AFM stiffness measurements of MDA^- (shTg2)^ + SF and MDA^- (shTg2)^ + MV^-^ primary tumors. (N=3+ tumors per condition; n = 128, 140). (d) Fraction of tumor area positive for αSMA (N=3). (e) Representative images of αSMA (red) and DAPI (blue) in MDA^- (shTg2)^ + SF and MDA^- (shTg2)^ + MV^-^ primary tumors. (f) Immunohistochemical staining of collagen and fibronectin in MDA^- (shTg2)^ + SF and MDA^- (shTg2)^ + MV^-^ primary tumors. (g) Representative BLI of MDA^- (shTg2)^ metastasis. (h) Representative image of macroscopic liver nodules. (i) Anti-GFP immunohistochemical staining of lungs and liver from MDA^- (shTg2)^ + SF and MDA^- (shTg2)^ + MV^-^ mice. (j) Quantification of percentage of lung tissue area GFP-po≡itive. (N=6, 8). (k) Quantification of percentage of liver tissue area GFP-po≡itive. (N=6, 8). (l) Illustration overviewing the mechanism of MDA^-^ metastasis. Data shown as mean +/− SEM.

### Clinical implications of Tg2 expression on breast cancer progression

Given that MV-Tg2 facilitate weakly migratory cancer cell metastasis, we investigated the clinical ramifications of these findings by examining Tg2 expression as a function of patient prognosis. Using the TNMplot database, we found that Tg2 expression of breast cancer patients increased from normal tissue to breast tumor tissue to metastatic tissue(47), indicating that Tg2 expression was correlated with breast cancer metastasis (Fig. 7a). Additionally, while distant metastasis free survival of breast cancer patients was weakly correlated with Tg2 expression (Fig. 7b), patients with triple negative breast cancer (TNBC) exhibited significantly decreased distant metastasis free survival with high Tg2 expression compared to low Tg2 expression(48) (Fig. 7c). Together, these findings further implicate Tg2 as an indicator of breast cancer progression and identify Tg2 as an important therapeutic target to prevent metastasis.

**Figure 7.**
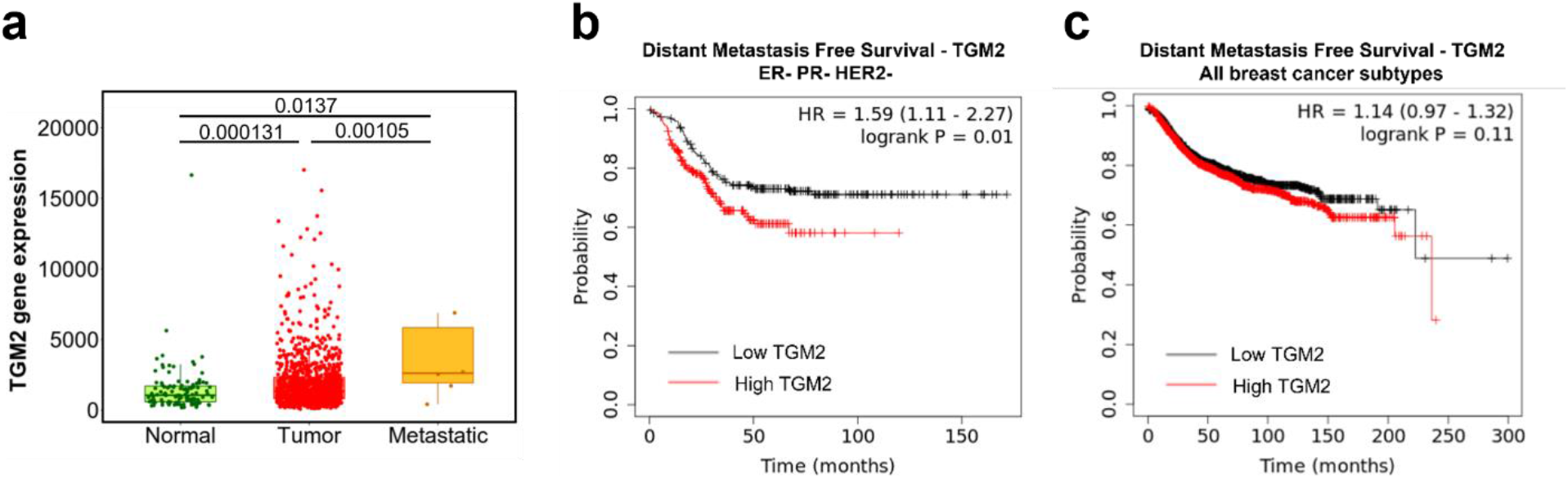
Clinical implications of Tg2 expression on breast cancer progression. (a) TGM2 gene expression, measured with RNA sequencing, of normal, tumor, and metastatic tissue from breast invasive carcinoma patients. (N=242, 7569, 82). Data adapted from TNMplot database. (b) Distant metastasis free survival Kaplan-Meier plot for TGM2 expression in all breast cancer subtypes. Data adapted from the Kaplan-Meier Plotter database. (c) Distant metastasis free survival Kaplan-Meier plot for TGM2 expression in ER, PR, and HER2 negative breast cancer. Data adapted from the Kaplan-Meier Plotter database.

## Discussion

Identifying the mechanisms of cancer cell dissemination away from the primary tumor is essential to develop new therapies to target metastatic cancer cells. Specifically, we show that in breast cancer, while highly migratory cells are capable of independent migration, weakly migratory cells release Tg2-rich MVs which activate fibroblasts which subsequently promote cancer cell dissemination and ultimately metastasis. Our data suggests that the interaction between weakly migratory cancer cells and stromal fibroblasts via MVs facilitates weakly migratory cancer cell dissemination and metastasis.

Our data supports the observation that cells do not need to be inherently migratory to metastasize(4,22,23,25–27). Others have shown that MDA-MB-231 isolated from bone metastases were less migratory than their primary tumor counterparts(22) and that highly metastatic subpopulations of MDA-MB-231 selected through repeated *in vivo* isolation of metastases were weakly migratory(23,25). Our work is the first to describe a mechanism by which cancer cells with a weak migration phenotype can manipulate cells within the tumor microenvironment to escape the primary tumor. Given the current emphasis on cell migration in cancer metastasis research, these findings highlight that cancer cell migration alone is not a sufficient indicator of metastatic potential. A focus on MV-mediated cell-cell signaling between cancer and stromal cells is crucial to fully understand the dynamics of cancer cell dissemination.

Our findings suggest a feedback loop between MV signaling, matrix remodeling, and fibroblast activation may exist. We previously reported that MV-mediated fibroblast activation was enhanced in high stiffness matrices mimicking the breast tumor periphery(38). Here, we show that Tg2-rich MVs stiffen the primary tumor, induce matrix deposition by stromal cells, activate fibroblasts, and promote cancer metastasis. Together, these data indicate that MV-induced fibroblast activation leads to increased tumor stiffening through matrix remodeling and this increased stiffness may feedback to prime fibroblasts for continued activation to further enhance cancer metastasis. Additionally, previous work revealed that transformation of normal fibroblasts *in vivo* by MDA-MB-231 MVs was dependent upon Tg2(38). Our results point to a role for these MV-Tg2-activated fibroblasts in mediating cancer cell escape from the primary tumor through matrix remodeling. As knockdown of Tg2 in MDA-MB-231 is known to decrease metastasis(49), we suspect this may due, in part, to reduced MV-mediated fibroblast activation and decreased cancer cell escape from the primary tumor.

Importantly, our work is the first to show that MV cargo varies based on cell phenotype. While it is known cancer cell phenotype affects the numbers of MVs released(50), our work reveals that differences in cancer cell phenotype within a single cell line actually correlates with differences in MV cargo. Additionally, while previous work showed that highly migratory cancer cells package migration-promoting cargo into EVs(51), our results suggest that heterogeneity in MV cargo based on migratory phenotype has downstream effects on fibroblast activation, tumor microenvironment remodeling, and ultimately metastasis. Our findings specifically show that Tg2 is not uniformly expressed by all breast cancer-derived MVs but rather dependent upon the migratory phenotype of the MV releasing cell, with increased Tg2 expression in MVs released by the weakly migratory breast cancer cells. Interestingly, the weakly migratory cells express high levels of Tg2 and E-cadherin(4). While Tg2 expression is generally linked with EMT and positively correlated with cancer cell invasion and migration(52–54) and E-cadherin is a marker of epithelial phenotypes and decreased cancer cell migration(24,26,55), Tg2 was recently identified as a marker of epithelial-mesenchymal plasticity and was found to be upregulated in cancer cells undergoing EMT only after a reversion to a secondary epithelial state(46). Taken together, these findings reveal that MV cargo changes as cells plastically change their EMT phenotype during the metastasis.

Additionally, our findings suggest that not all MVs released from a cell population will homogenously communicate with surrounding cells. In cancer, targeting MV populations that contain cargo (such as Tg2) to remodel the tumor microenvironment may be crucial to reduce metastatic potential. Importantly, other groups have shown that exosomes affect later stages of cancer metastasis, including angiogenesis and pre-metastatic niche formation(46,56,57). As exosomes and MVs contain many of the same cancer-promoting signaling proteins, such as Tg2, VEGF, and TGFβ(46,58–61), MVs may also signal to endothelial cells to promote angiogenesis and travel to secondary sites to promote pre-metastatic niche formation. Taken together, these findings reveal that identifying the MV populations and MV cargo capable of promoting progression through the metastatic cascade is essential to prevent metastasis and highlight MVs as a potential target for new anti-metastatic cancer therapies.

In summary, the relationship between cancer cell migratory ability and dissemination away from the primary tumor is complex. While highly migratory cells are capable of independent migration, our work identifies a population of weakly migratory, highly metastatic breast cancer cells which escape the primary tumor via MV-Tg2-mediated fibroblast activation. As we further define the relationship between cancer cell migration and metastatic potential and the consequences of MV signaling on cancer progression, these findings are likely to have broader implications in designing modalities and therapies to detect and target metastatic cancer cells.

## Materials and Methods

### Cell culture and reagents

MDA-MB-231 malignant mammary adenocarcinoma cells (HTB-26, ATCC, Manassas, VA), NIH 3T3 fibroblasts, and all modified cell lines were maintained in Dulbecco’s Modified Eagle’s Media (DMEM) (ThermoFisher Scientific, Waltham, PA) supplemented with 10% fetal bovine serum (Atlanta Biologicals, Flowery Branch, GA) and 1% penicillin-streptomycin (ThermoFisher Scientific). All cells were cultured at 37°C and 5% CO_2_.

Primary antibodies used were rabbit anti-flotillin-2 (3436; Cell Signaling Technology, Danvers, MA), mouse anti-α smooth muscle actin (M0851, DAKO, Santa Clara, CA), mouse anti-beta actin (A5316, Millipore Sigma, Burlington, MA), mouse anti-tissue transglutaminase 2 (ab2386, Abcam, Cambridge, UK), goat anti-fibronectin (sc6953, Santa Cruz, Dallas, TX), rabbit anti-fibronectin (F3648, Millipore Sigma), rabbit anti-focal adhesion kinase (3285, Cell Signaling Technology, Danvers, MA), rabbit anti p-focal adhesion kinase (Tyr397) (3283, Cell Signaling Technology), and mouse-anti GAPDH (MAB374, Millipore Sigma). Secondary antibodies used were HRP anti-rabbit (Rockland, Limerick, PA), HRP anti-mouse (Rockland), AlexaFluor 488 conjugated to donkey anti-goat (Life Technologies, Carlsbad, CA), AlexaFluor 488 conjugated to donkey anti-rabbit (Life Technologies), AlexaFluor 488 conjugated to donkey anti-mouse (Life Technologies), and AlexaFluor 568 conjugated to donkey anti-mouse (Life Technologies). Actin was stained using Texas Red Phalloidin (Life Technologies).

### Phenotypic sorting of MDA-MB-231 breast cancer cells

MDA-MB-231 breast cancer cells were phenotypically sorted based on their ability to migrate through a collagen gel on top of a Transwell insert as previously described(4). Briefly, cancer cells were seeded on a thin layer of 1 mg/ml collagen (Corning, Corning, NY) on top of a Transwell insert with 8 μm pores (Greiner Bio-One, Kremsmunster, Austria). A serum gradient was applied and cancer cells were allowed to migrate for four days. After four days, highly migratory and weakly migratory cells were collected and reseeded in fresh Transwells. After 20 rounds of purification, cells that repeatedly migrated through the assay were termed ‘highly migratory’ (MDA^+^) and cells that never migrated through the assay were termed ‘weakly migratory’ (MDA^-^).

### Modified cell lines

MDA^+^ and MDA^-^ were stably transduced with either FUW-GFP-E2A-fluc or FUW-mCherry-E2A-rluc, both created in-house. NIH 3T3 fibroblasts were transduced with either Life-Act eGFP (#84383, Addgene, Watertown, MA) or FUW-mCherry-E2A-rluc. To generate a Tg2-knockdown and a scrambled control cell line, MDA^- (GFP/luc)^ were subjected to lentiviral transduction with either the Tg2-targeting MISSION shRNA plasmid (SHCLND-NM_004613: TRCN0000272816) (MDA^- (scr)^) or the MISSION scr.1-puro scrambled control plasmid DNA (SHC001; Millipore Sigma) (MDA^- (shTg2)^). To generate a Tg2 overexpressing cell line, MDA^+^ were stably transduced with an FUW-Tg2 plasmid created in-house.

### Orthotopic mammary metastasis mouse model

Orthoptic injection of NOD SCID gamma (NSG) mice was conducted as previously described(4). Briefly, female NSG mice, 6-8 weeks of age, were injected subcutaneously at the fourth mammary fat pad with either MDA^+^, MDA^-^, MDA^- (scr)^, or MDA^- (shTg2)^, lentivirally transduced with GFP and firefly luciferase tags. In mouse experiments with MVs, either serum-free media alone or approximately 1*10^7^ MV^-^ suspended in serum-free media were injected subcutaneously at the primary tumor site every three days from one to six weeks post cancer cell injection. For bioluminescent imaging, mice were injected with 30 mg/mL D-luciferin (Gold-Bio, St Louis, MO) and imaged weekly on an IVIS Lumina III Series (Caliper LifeSciences, Hopkinton, MA). After 4-6 weeks, primary tumors were removed using sterile surgical technique. Primary tumor size was measured with calipers. Primary tumors were cut in half and either snap-frozen in dry ice or fixed with 4% v/v paraformaldehyde (Electron Microscopy Sciences, Hartfield, PA). At the endpoint, mouse lung and liver were collected and fixed in 4% v/v paraformaldehyde for 24 hours and sent to the Vanderbilt Tissue Pathology Shared Resource for paraffin embedding and sectioning.

### Atomic force microscopy of primary mouse tumors

Primary tumor samples cryopreserved in O.C.T compound were cut into 20 μm sections. Prior to AFM stiffness measurements, samples were thawed at room temperature for 3 minutes, and maintained in 1X Halt Protease Inhibitor Cocktail (78438, ThermoFisher Scientific). A thermoplastic coverslip (ThermoFisher Scientific) with a 6 μm biopsy punch hole was superglued on top of the tumor section. An ImmEdge hydrophobic pen (Vector Laboratories, Burlingame, CA) was used to draw a small circle around the biopsy punch hole. AFM measurements were performed using a MFP3D-BIO inverted optical AFM (Asylum Research, Santa Barbara, CA), with an inverted fluorescent Zeiss Observer Z.1 microscope with a 10x/0.3 N.A. objective. A silicon nitride cantilever with a 5 μm diameter spherical borosilicate glass tip and a spring constant of 0.06 N/m (Novascan Technology, Boone, IA) was used. Samples were indented at a 2 μm/second loading rate, with a maximum indentation force of 5 nN. To obtain tumor stiffness measurements, IGOR PRO Software (Asylum Research) was used. At least two force maps of each primary tumor sample were obtained. AFM data was fit to a Hertz model with a Poisson’s ratio of 0.5.

### qPCR of tumors

Snap frozen primary tumors were prepared for RNA isolation by TRIZOL digestion and subsequent homogenization using a TissueLyser II (Qiagen, Hilden, Germany) with a 5 mm stainless steel bead for 2 min at 30 Hz. After homogenization, samples were incubated at room temperature for 5 minutes in TRIZOL. Chloroform was added to the homogenized sample and incubated at room temperature for 3 minutes. Samples were centrifuged at 10,000 x g for 30 minutes at 4C. The upper aqueous phase was separated and mixed with 70% ethanol. RNA was subsequently isolated using the RNeasy Mini Kit (Qiagen). DNA was synthesized from RNA using the iScript cDNA Synthesis Kit (Bio-Rad Laboratories, Hercules, CA). qPCR was performed with the iQ SYBR Green Supermix (Bio-Rad Laboratories) per manufacturer protocols. The primer sequences for each gene are listed in Supplementary Table 1.

**Supplementary Table 1.**
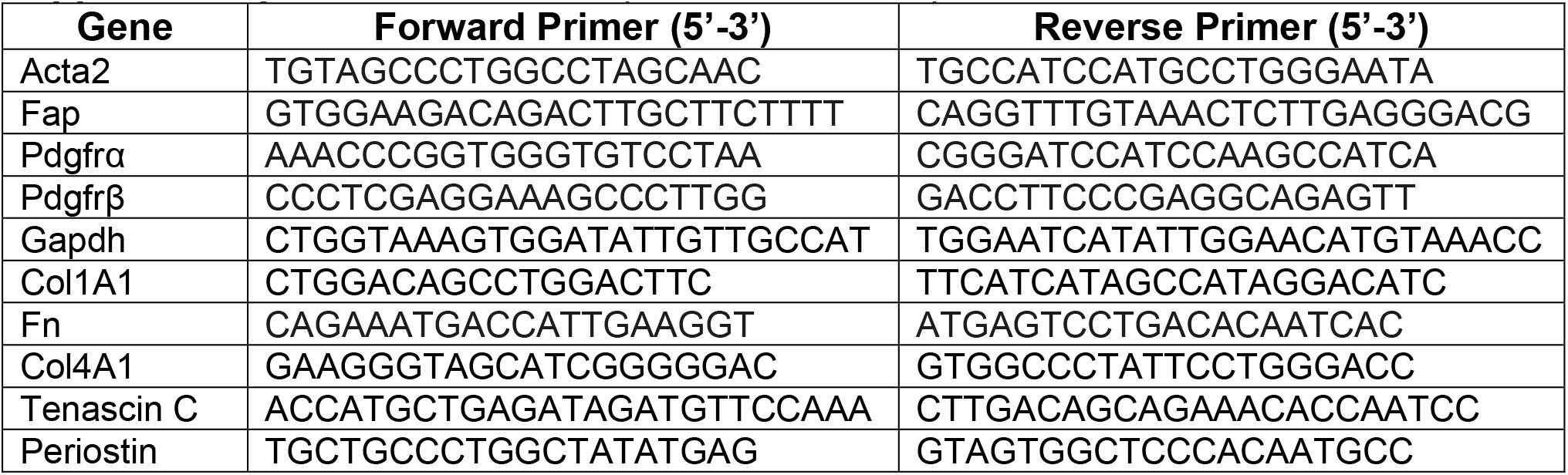
Primer sequences used for quantitative real-time PCR.

### Immunohistochemical staining of mouse tissues

Primary tumors were fixed, paraffin-embedded, and mounted into 5 μm thick sections at the Vanderbilt Tissue Pathology Shared Resource. For collagen staining, tumors were stained using the picrosirius red stain kit (Polysciences Inc., Warrington, PA). For fibronectin staining, tumors were stained using the Abcam IHC-Paraffin Protocol. Briefly, 5 μm thick sections of primary tumors were deparaffinized and rehydrated using a series of washes of xylene, ethanol, and water. Heat-induced epitope retrieval was completed using a sodium citrate buffer. Tumor sections were blocked for 2 hours in tris buffered saline (TBS) + 10% fetal bovine serum + 1% bovine serum albumin at room temperature. Samples were incubated overnight in rabbit anti-fibronectin (1:500) in TBS + 1% bovine serum albumin. After washing, samples were incubated in TBS + 0.3% hydrogen peroxide for 15 minutes to suppress endogenous peroxidase activity. Samples were incubated in HRP anti-rabbit (1:200) for 1 hour. Staining was developed using the DAB chromogen (8059S, Cell Signaling Technology) and samples were subsequently counterstained using Mayer’s hematoxylin (Millipore Sigma). Tumor samples were then dehydrated, cleared, and mounted. Anti-GFP staining of liver and lungs was completed by the Vanderbilt Tissue Pathology Shared Resource. Whole slide imaging and quantification of immunostaining were performed in the Digital Histology Shared Resource at Vanderbilt University Medical Center (www.mc.vanderbilt.edu/dhsr).

### Immunofluorescent staining of mouse tissues

Primary tumors were frozen and cut into 20 μm thick sections at the Vanderbilt Tissue Pathology Shared Resource. Tumor sections were thawed, fixed for 10 minutes in 4% v/v paraformaldehyde, and permeabilized in 1% Triton X-100 (Millipore Sigma). Sections were blocked for 2 hours in PBS + 10% fetal bovine serum + 5% donkey serum + 5% goat serum. Samples were incubated overnight in mouse anti-αSMA (1:200). After washing, samples were incubated in AlexaFluor 568 conjugated to donkey anti-mouse (1:200) and DAPI (1:500) for 2 hours. Sections were mounted for imaging using Vectashield Mounting Media (Vector Laboratories). Sections were imaged with a 10x/0.3 N.A. objective on a Zeiss LSM800 confocal laser-scanning microscope.

### MV isolation and characterization

MDA^+^, MDA^-^, MDA^- (scr)^, MDA^- (shTg2)^, and MDA^+^ (^Tg2-FUW^) were incubated overnight in serum-free DMEM. The conditioned media was removed from the cells and centrifuged at 400 RPM. The supernatant was removed and again centrifuged at 400 RPM. The medium was then filtered through a 0.22 μm SteriFlip filter unit (Millipore Sigma) and rinsed with serum-free media. The MVs retained by the filter were resuspended in serum-free media. Nanoparticle tracking analysis (ZetaView ParticleMetrix, Germany) was used to determine the size and number of isolated MVs from MDA^+^ (MV^+^), MDA^-^ (MV^-^), MDA^- (scr^) (MV^- (scr)^), and MDA^- (shTg2)^ (MV^- shTg2)^).

### Western blotting

Isolated MVs were rinsed with PBS on a 0.22 μm SteriFlip filter unit and lysed with Laemmli buffer. MDA^+^, MDA^-^, and NIH 3T3 fibroblasts were lysed with Laemmli buffer. For T101 experiments, MV^-^ were incubated for 30 minutes with either DMSO or 10 μM T101. MVs were re-filtered and resuspended in serum-free media. NIH 3T3 fibroblasts were cultured with either MV^- (DMSO)^ or MV^- (T101)^ for 24 hours before lysis with Laemmli buffer. All lysates were resolved by SDS-PAGE and transferred to PVDF membranes. Transferred membranes were blocked with either 5% milk or 5% bovine serum albumin in TBS-Tween. Membranes were incubated overnight in mouse anti-α smooth muscle actin (1:1000), rabbit anti-flotillin-2 (1:1000), mouse anti-tissue transglutaminase 2 (1:1000), rabbit anti-focal adhesion kinase (1:1000), rabbit anti-phospho focal adhesion kinase (1:1000), and mouse anti-GAPDH (1:2000) at 4°C. Membranes were then incubated in horseradish peroxidase-conjugated secondary antibody (1:2000) in 5% milk or 5% bovine serum albumin in TBS-Tween for 1 hour at room temperature. Samples were imaged with an Odyssey Fc (LI-COR Biosciences, Lincoln, NE) after the addition of SuperSignal West Pico or West Dura Chemiluminescent Substrates (ThermoFisher Scientific).

### Polyacrylamide gel preparation

Polyacrylamide (PA) gels were fabricated as described elsewhere(62). Briefly, the ratio of acrylamide (40% w/v; Bio-Rad, Hercules, CA) to bis-acrylamide (2% w/v; Bio-Rad) was varied to tune gel stiffness to 20 kPa, previously shown to enhance fibroblast response to MVs(38). 20 kPa PA gels were fabricated using an acrylamide: bis-acrylamide ratio of 12%:0.19%. PA gels were coated with 0.1 mg/ml rat tail type I collagen (Corning).

### Fibronectin immunofluorescence and analysis

NIH 3T3 fibroblasts were seeded on 20 kPa PA gels in 1.6 mL of DMEM + 1% FBS. Cells were treated with either 400 μL of serum-free media or approximately 3*10^7^ MVs suspended in 400 μL serum-free media for 24 hours. Cells were first fixed with 3.2% v/v paraformaldehyde and subsequently permeabilized with a 3:7 ratio of methanol to acetone. Cells were incubated with goat anti-fibronectin (1:100) overnight. The cells were washed and incubated with AlexFluor 488 conjugated to donkey anti-goat (1:200), TexasRed phalloidin (1:500), and DAPI (1:500) for 1 hour. To image, gels were inverted onto a drop of Vectashield Mounting Media placed on a glass slide. Fluorescent images were acquired with a 20x/1.0 N.A. water-immersion objective on a Zeiss LSM700 Upright laser-scanning microscope. Cells stained with phalloidin were outlined in ImageJ to calculate cell area. Area of fibronectin was measured by using the threshold function in ImageJ to calculate area of positive stain. The ratio of fibronectin area to cell area was calculated.

### Phalloidin and αSMA immunofluorescence and analysis

NIH 3T3 fibroblasts were seeded on 20 kPa PA gels in 1.6 mL of DMEM + 1% FBS. Cell media was additionally supplemented with either 400 μL of serum-free media or approximately 3*10^7^ MVs suspended in 400 μL serum-free media. After 24 hours, cells were fixed with 3.2% v/v paraformaldehyde and permeabilized with 1% Triton X-100. Cells were blocked with 3% bovine serum albumin in 0.02% Tween in PBS and then incubated for 3 hours at room temperature with mouse anti-alpha smooth muscle actin (1:100). After being washed, cells were incubated for 1 hour with AlexaFluor 488 conjugated to donkey anti-mouse (1:200). The cells were washed and F-actin and nuclei were stained with TexasRed phalloidin (1:500) and DAPI (1:500), respectively. To image, gels were inverted onto a drop of Vectashield Mounting Media placed on a glass slide. Fluorescent images were acquired with a 20x/1.0 N.A. water-immersion objective on a Zeiss LSM700 Upright laser-scanning microscope. For αSMA expression, cells stained with phalloidin were outlined in ImageJ. Cell area was overlaid onto αSMA images and integrated density was measured. Corrected total cell αSMA fluorescence (CTCF) was calculated by subtracting the cell area multiplied by the mean fluorescence of the background by the integrated density of the cell.

### EdU proliferation assay

NIH 3T3 fibroblasts were serum-starved for 6 hours and subsequently seeded on 20 kPa PA gels in 1.6 mL of DMEM + 1 % FBS. Cell media was additionally supplemented with either 400 μL of serum-free media or approximately 3*10^7^ MVs suspended in 400 μL serum-free media. After 24 hours, 10 μM 5-ethynyl-2’-dexoyuridine (EdU, ThermoFisher Scientific) was added to the culture media for 2 hours. Cells were fixed with 3.2% v/v paraformaldehyde and stained with the Click-iT EdU Kit (ThermoFisher Scientific) following the manufacturer’s instructions. Nuclei were counterstained with DAPI (1:500). Cells were imaged with a 20x/1.0 N.A. water-immersion objective on a Zeiss LSM700 Upright laser-scanning microscope. The percentage of EdU incorporation was calculated as the ratio of EdU positive cells to the total number of cells.

### Traction force microscopy

Traction force microscopy was performed as previously described(63). Briefly, 20 kPa PA gels, embedded with 0.5 μm diameter fluorescent beads (ThermoFisher Scientific), were prepared. NIH 3T3 fibroblasts were allowed to adhere for 24 hours in 1.6 mL of DMEM + 1% FBS supplemented with either 400 μL of serum-free media or approximately 3*10^7^ MVs suspended in 400 μL serum-free media. After 24 hours, phase contrast images of fibroblasts and fluorescent images of the beads at the surface of the PA gel were acquired. Fibroblasts were removed from the PA gel using 0.25% trypsin/EDTA (Life Technologies) and a fluorescent image of the beads was acquired after cell removal. Bead displacements between the stressed and null states and fibroblast area were calculated and analyzed using the LIBTRC library developed by M. Dembo (Dept. of Biomedical Engineering, Boston University)(64). Outliers were removed using the ROUT method with Q = 0.2%.

### Spheroids

MDA^+^, MDA^-^, MDA^- (scr)^, MDA^- (shTg2)^ and NIH 3T3 fibroblasts, transduced with either GFP or mCherry, were resuspended in spheroid compaction media containing 0.25% methylcellulose (STEMCELL, Vancouver, Canada), 4.5% horse serum (ThermoFisher Scientific), 18 ng/mL EGF (ThermoFisher Scientific), 90 ng/mL cholera toxin (Millipore Sigma), 90 U/mL penicillin (Life Technologies), and 90 ug/mL streptomycin (Life Technologies) in DMEM/F12 (ThermoFisher Scientific). A 2:1 ratio of cancers cells to fibroblasts were added to wells of a round-bottom 96 well plate to generate 5,500 cell spheroids. The plate was centrifuged at 1100 rpm for 5 minutes at room temperature and subsequently incubated at 37 degrees for 72 hours to allow for spheroid compaction.

After 72 hours of spheroid compaction, spheroids were embedded into 4.5 mg/mL collagen gels. Briefly, 4.5 mg/mL collagen gels were generated by mixing 10 mg/ml stock collagen (Rockland), 0.1% acetic acid, culture media, 10x HEPES (Millipore Sigma), and 1 N NaOH (Millipore Sigma) and added to wells of a 24 well plate. Compacted spheroids were placed into the middle of the collagen gel without touching the bottom of the well plate. The embedded spheroids were placed in a 37°C incubator for 10 minutes to polymerize, then the plate was flipped upside-down for another 30 minutes to avoid spheroid adhesion on the bottom of the well. Spheroids were rehydrated every 24 hours with 300 μL of 1% DMEM and supplemented with either 200 μL of serum-free DMEM or 1.5*10^7^ MVs suspended in 200 μL serum-free DMEM. Spheroids were imaged every 24 hours for 48 hours using a Zeiss LSM800 inverted confocal microscope equipped with an environmental control chamber. Spheroid images were captured using a 10X/0.3 NA objective. The projected spheroid area, the spheroid diameter, and the maximum cancer cell migration distance from the spheroid core was measured. The expansion index (A_i_/A_0_-1) was calculated to quantify expansion 48 hours post spheroid embedding.

### iTRAQ Proteomics and Analysis

After MV isolation, MV^+^ and MV^-^ were lysed in a buffer composed of 2% Nonidet P-40 (Millipore Sigma), 0.5% sodium deoxycholate (Millipore Sigma), 300 mM sodium chloride (Millipore Sigma), and 50 mM Tris pH 8. Lysates were incubated for 30 minutes at 4°C and centrifuged at 14,000 x g for 15 minutes at 4°C to pellet non-solubilized proteins. iTRAQ proteomics of lysates was completed by the Vanderbilt Mass Spectrometry Research Center Proteomics Core. Briefly, enzyme digestion of lysates was used to generate proteolytic peptides. MV^+^ and MV^-^ peptides were labeled with 117 and 115 iTRAQ reagents, respectively. Samples were subsequently mixed, fractioned using liquid chromatography, and analyzed via tandem mass spectrometry. A database search using the fragmentation data identified the labeled peptides and their corresponding proteins. Protein set enrichment analysis was completed using the PSEA-Quant algorithm(45). The REVIGO web tool was used to remove redundant GO terms(65).

### Statistical analysis

All statistical analysis was performed using GraphPad Prism 7 (GraphPad Software, La Jolla, CA) or Excel 2016 (Microsoft, Redmond, WA). Where appropriate, data were compared with a student’s t-test or with a two-way analysis of variance (ANOVA) with Sidak multiple comparisons test. All data is reported as mean ± standard error (SE) unless otherwise notated.

## Acknowledgements

This work was supported by the W.M. Keck Foundation, the National Institutes of Health (GM13117) to C.A.R.-K. This work was also supported by National Science Foundation Graduate Research Fellowship Awards under Grant No. 1937963 to S.C.S. and J.A.M and Grant No. DGE-1650441 to L.A.H. and the Scholarship for the Next Generation of Scientists from the Cancer Research Society, the National Cancer Institute Grant K99CA212270, and the Canada Research Chair program awarded to F.B. We thank Alissa Weaver for the use of the ZetaView ParticleMetrix. We acknowledge the Tissue Pathology Shared Resource supported by NCI/NIH Cancer Center Support Grant 5P30 CA68485-19, the Vanderbilt Mouse Metabolic Phenotyping Center Grant 2 U24 DK059637-16, and the Shared Instrumentation Grant S10 OD023475-01A1 for the Leica Bond Rx.

## Author Contributions

S.C. Schwager, L.A. Hapach, F. Bordeleau, M.A. Antonyak, R.A. Cerione, and C.A. Reinhart-King designed research; S.C. Schwager, L.A. Hapach, C. Carlson, Y.E. Chu, J.A. Mosier, W. Wang performed experiments; S.C. Schwager, L.A. Hapach, C. Carlson, Y.E. Chu, A.L. Jayathilake, and T. McArdle analyzed data; L.A. Hapach, M.A. Antonyak, R.A. Cerione, and C.A. Reinhart-King aided in experimental design; and all authors contributed to writing the report.

## Supplemental Figures

**Supplemental Figure 1.**
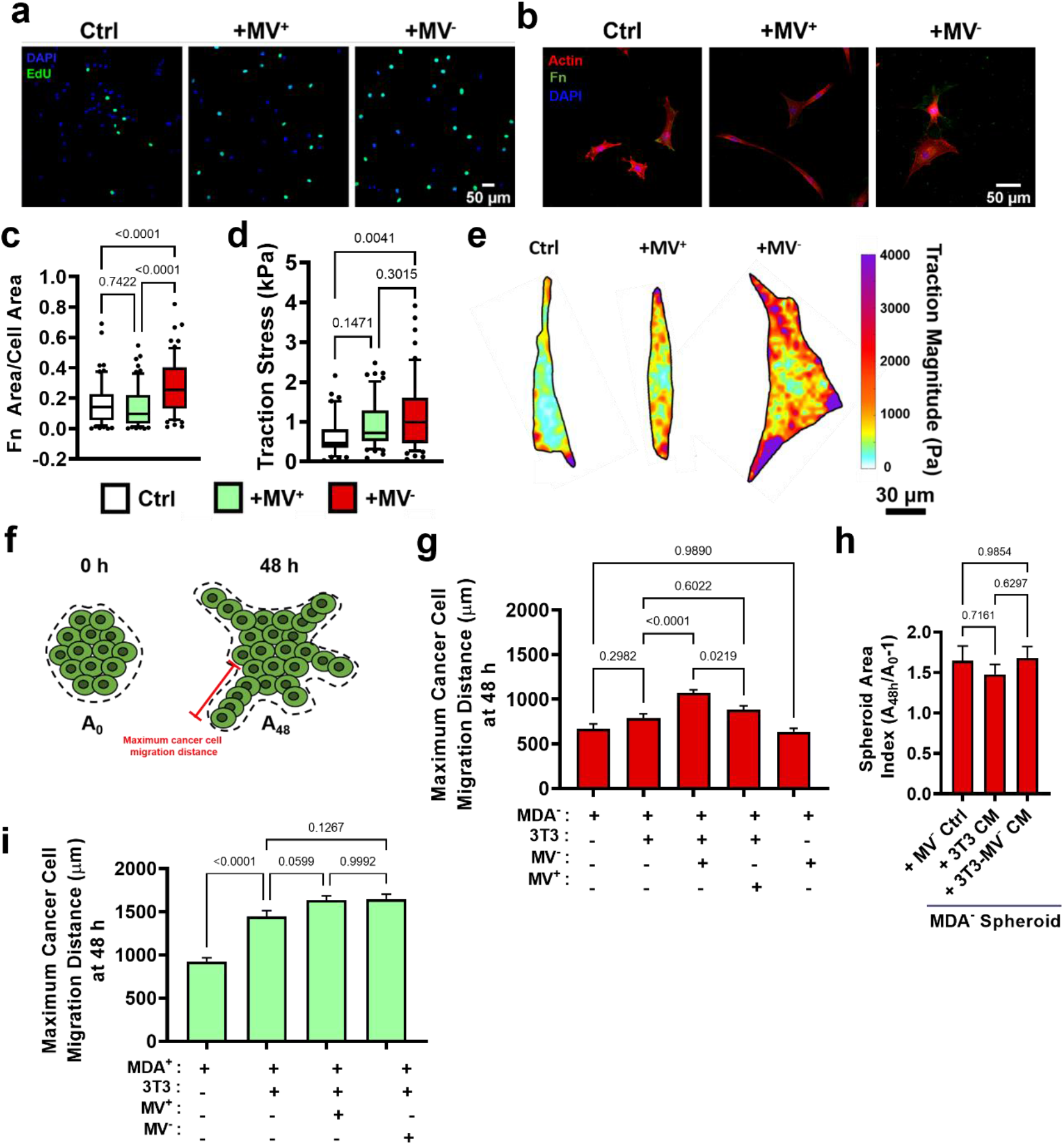
(a) Representative images of fibroblast EdU incorporation after culture in control conditions (Ctrl), with MV^+^ (+MV^+^), or with MV^-^ (+MV^-^). DAPI (blue); EdU (green). (b) Representative images of fibroblast fibronectin deposition after culture in Ctrl, +MV^+^, or +MV^-^ conditions. Actin (red); fibronectin (green); DAPI (blue). (c) To ensure that the increased fibronectin area was not a result of increased fibroblast spreading, fibronectin area was normalized to cell area. (N=3; n=66, 66, 61) (d) Fibroblast traction stress (calculated as fibroblast traction force / cell area) after culture in Ctrl, +MV^+^, or +MV^-^ conditions. (N=3; n=41, 57, 58). (e) Representative fibroblast traction stress maps. (f) Schematic of spheroid outgrowth measurements. (g) Maximum distance of cancer cell migration away from spheroid core 48 hours post spheroid (i) Maximum distance of cancer cell migration away from spheroid core 48 hours post spheroid embedding. (N=3+; n=28, 32, 54, 24). Bar graphs show mean +/− SEM. Box and whisker plots show median and 25^th^-75^th^ (box) and 10^th^-90^th^ (whiskers) percentiles. P-values determined using a one-way ANOVA with Tukey’s test for multiple comparisons.

**Supplemental Figure 2.**
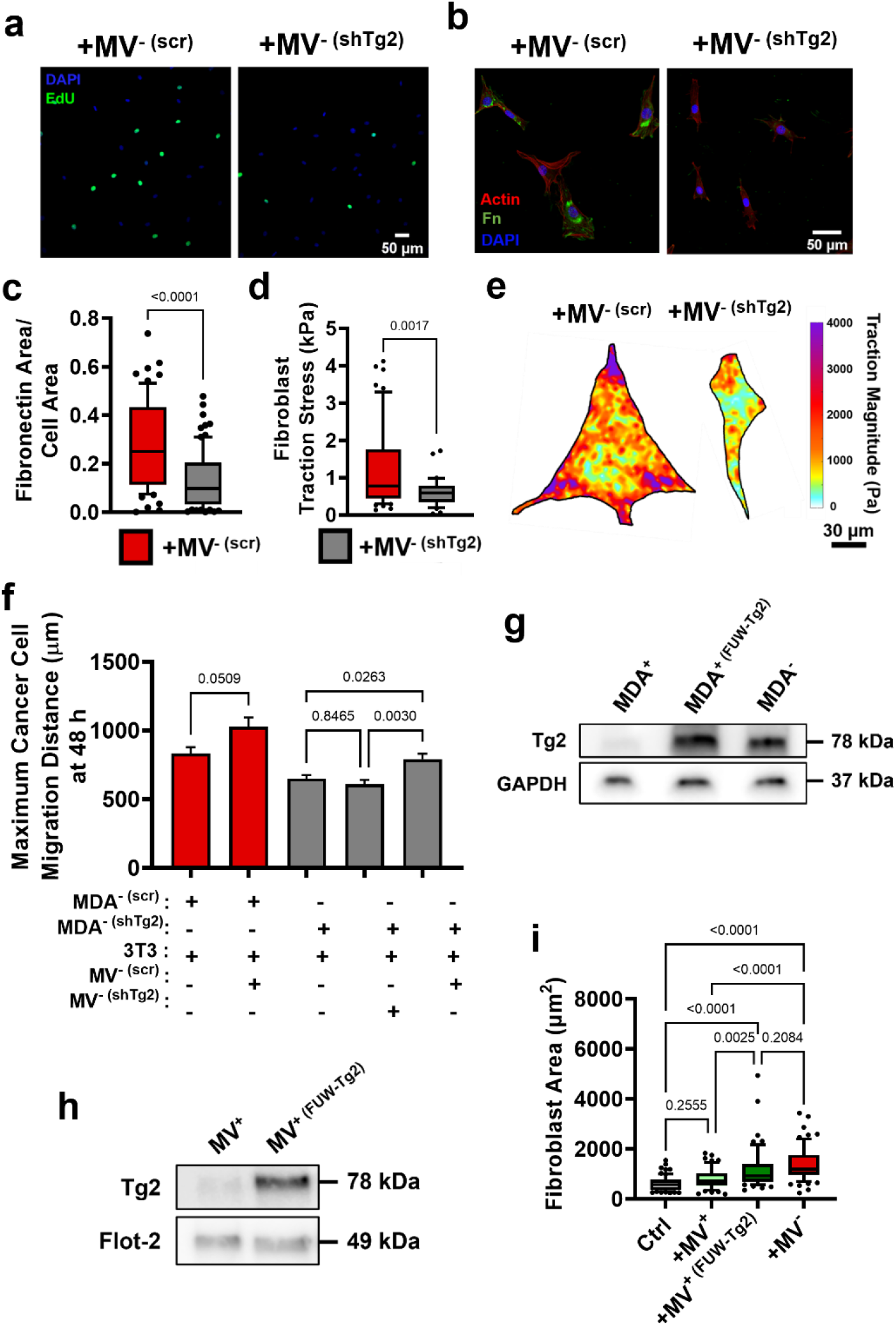
(a) Representative images of fibroblast EdU incorporation after culture with MV^- (scr)^ (+MV^- (scr)^) or MV^- (shTg2)^ (+MV^- (shTg2)^). DAPI (blue); EdU (green). (b) Representative images of fibroblast fibronectin deposition after culture in +MV^- (scr)^ or +MV^- (shTg2)^ conditions. Actin (red); fibronectin (green); DAPI (blue). (c) Fibronectin area normalized to fibroblast cell area. (N=3; n=62, 82). (d) Fibroblast traction stress (calculated as fibroblast traction force / cell area) after culture in +MV^- (scr)^ or +MV^- (shTg2)^ conditions. (N=3; n=56, 33). (e) Representative fibroblast traction stress maps. (f) Maximum distance of cancer cell migration away from spheroid core 48 hours post spheroid embedding. (N=3+; n= 23, 23, 83, 60, 37). (g) Western blot for Tg2 and GAPDH in MDA^+^, MDA^+ (FUW-Tg2)^, and MDA^-^. (h) Western blot for Tg2, Flotillin-2, and B-actin in MV^+^ and MV^+ (FUW-Tg2)^. (i) Fibroblast cell area after culture in control conditions (Ctrl), with MV^+^ (+MV^+^), with MV^+ (FUW-g2)^ (+MV^+ (FUW-Tg2)^, or with MV^-^ (+MV^-^). (N=3; n=70, 57, 56, 64). Bar graphs show mean +/− SEM. Box and whisker plots show median and 25^th^-75^th^ (box) and 10^th^-90^th^ (whiskers) percentiles. P-values determined using an unpaired Student’s t-test or a one-way ANOVA with Tukey’s test for multiple comparisons.

